# *Yuel*: Compound-Protein Interaction Prediction with High Generalizability

**DOI:** 10.1101/2021.07.06.451043

**Authors:** Jian Wang, Nikolay V. Dokholyan

## Abstract

Virtual drug screening has the potential to revolutionize the stagnant drug discovery field due to its low cost and fast speed. Predicting binding affinities between small molecules and the protein target is at the core of computational drug screening. Deep learning-based approaches have recently been adapted to predict binding affinities and claim to achieve high prediction accuracy in their tests, however, we show that current approaches are not reliable for virtual drug screening due to the lack of generalizability, i.e. the ability to predict interactions between unknown proteins and unknown small molecules. To address this shortcoming, we develop a compound-protein interaction predictor, Yuel. Upon comprehensive tests on various datasets, we find that out of all the deep-learning approaches surveyed, only Yuel can predict interactions between unknown compounds and unknown proteins. Additionally, Yuel can also be utilized to identify compound atoms and proteins residues that are involved in the binding.

## INTRODUCTION

Drug discovery^1^ has strongly benefited from virtual drug screening^2^ techniques based on either molecular docking^3–6^ of a library of compounds to a target protein, or cheminformatics-based approaches that utilize machine learning to derive knowledge from known binders to this target. Yet, molecular docking is usually time-consuming, while cheminformatics-based approaches^7^ suffer from low accuracy when no ligand for a given target is known. Recently, deep learning has been widely applied to various areas in biology and is seemingly heralding a trend to supersede traditional methods due to its unparalleled performance, e.g., AlphaFold2^8^ has stunned the protein structure communication by sweeping the biennial protein 3D structure prediction competition, CASP (the critical assessment of protein structure prediction)^9^. In the virtual drug screening field, several deep learning-based neural networks^10–13^ have emerged aiming at predicting compound-protein interaction (CPI), such as DeepDTA^10^, DeepConv-DTI^11^, DeepAffinity^12^, and MONN^13^. However, considering that the chemical space of potential drugs may be as large as 10^60^ small molecules^14^ and the human proteome consist of millions of protein species^15,16^, a natural question that arises is whether current neural network models are reliable for drug virtual screening when training on a tiny dataset: typical compound-protein affinity datasets, such as the Davis^17^ and the PDBbind^18^ datasets, only contain ten thousand to hundred thousand affinity data. Unfortunately, we evaluate current methods and find that most are merely memorizing the inputs and outputs, thus lacking the ability to predict the interaction between unknown compounds and unknown proteins, which we refer to as generalizability. Here, we develop a new compound-protein interaction predictor, *Yuel* (Figure 1A), which predicts CPI with high generalizability. Upon comprehensive tests on various datasets, Yuel is the only method that can predict interactions between unknown compounds and unknown proteins.

**Figure 1.**
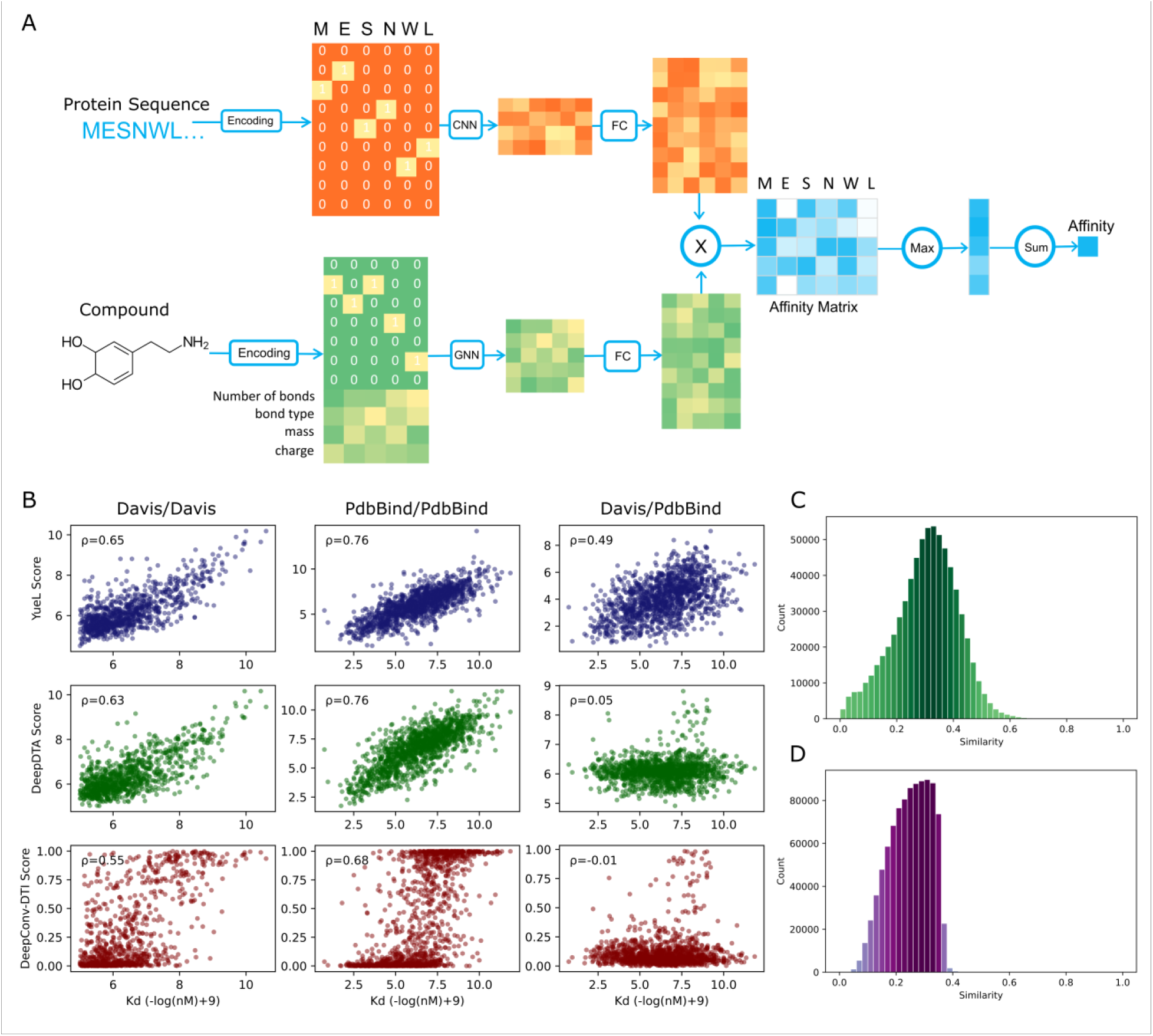
Test of the ability of Yuel to predict compound-protein interaction. (A) The architecture of Yuel. (B) Test of the performance of Yuel, DeepDTA, and DeepConv-DTI on the Davis and the PDBbind dataset. (C) The histogram of the similarities of compounds between the Davis dataset and the PDBbind dataset. (D) The histogram of the similarities of proteins between the Davis dataset and the PDBbind dataset. The average value of the similarities between compounds in the two datasets is 0.35, and the average similarity of proteins in the two dataset is 0.27.

## RESULTS & DISCUSSION

### Yuel predicts compound-protein interaction with high generalizability

We train and test Yuel and two other neural networks, DeepDTA^10^ and DeepConv-DTI^11^, on two datasets (PDBbind^18^ and Davis^17^). We do not compare Yuel to DeepDTA and DeepConv-DTI because MONN needs specific pairwise interactions between atoms in compounds and residues in proteins for training and DeepAffinity does not provide convenient utilities for training with custom datasets. We divide each of the two datasets into a training set and a test set at a ratio of 8 to 2, resulting in four datasets (Davis/train, Davis/test, PDBbind/train, PDBbind/test). Cross-validation is not performed because we will use independent datasets for testing. We first train Yuel, DeepDTA, and DeepConv-DTI on the training set (Davis/train and PDBbind/train) and test them on the corresponding test set (Davis/test and PDBbind/test). By training on Davis/train, the Spearman correlation between the experimental affinities and predicted affinities (Figure 1B) of Yuel, DeepDTA, and DeepConv-DTI on Davis/test are 0.65, 0.63, and 0.55, respectively; by training on PDBbind/train, the Spearman correlation on PDBbind/test are 0.76, 0.76, and 0.68, respectively. All three methods demonstrate high predictive values, with the performance of Yuel and DeepDTA demonstrating similar performance which is slightly higher than DeepConv-DTI. However, this significant result for all three neural networks is not surprising as the training and test sets are sourced from the same dataset, thus many proteins and compounds are shared between the training set and the test set.

To evaluate the neural networks’ ability to predict interactions between unknown proteins and unknown compounds, we train the models on Davis and test them on PDBBind. Compounds and proteins are dissimilar in these two datasets (Figure 1C&1D), so PDBbind can be used as an independent dataset when training on Davis. Further, Davis has far less proteins (379) and compounds (68) than PDBbind (proteins: 10251; compounds: 2509). We find the Spearman correlations of *Yuel*, DeepDTA, and DeepConv-DTI are 0.489, 0.054, and −0.015 respectively, suggesting that DeepDTA, and DeepConv-DTI cannot predict the interaction between unknown proteins and unknown compounds, while *Yuel* can. Therefore, Yuel can predict a large number of interactions between unknown proteins and unknown compounds by training on a small number of dissimilar proteins and dissimilar compounds, thus showing its high generalizability. We further train the three models on Davis and test them on a third dataset Metz^19^. The Metz dataset contains interactions of 172 different protein kinases and more than 3800 compounds. *Yuel* still outperforms DeepDTA and DeepConv-DTI in terms of the Spearman correlation (Figure S1).

### Yuel addresses the memorizing issue that exists in other neural network models

To test if networks are purely memorizing protein sequences or compound structures, we use PDBbind/train for training, but then shuffle the residues of each protein sequence in PDBbind/test for testing. Thus, the new protein sequences in the shuffled set are all chimeric and not supposed to bind to the compounds. However, we find that the shuffling does not significantly affect the performance of DeepDTA and DeepConv-DTI (Spearman correlation is 0.7 and 0.6, respectively) (Figure 2). On the contrary, *Yuel’*s predictions fall drastically to a Spearman correlation of 0.2 (Figure 2A), in line with expectations. Thus, DeepDTA and DeepConv-DTI are essentially predicting CPI by memorizing the compound structures. The memorization characteristic of DeepDTA and DeepConv-DTI may be due to the low completeness (see Methods) of the PDBbind dataset: each affinity data is nearly distinctly corresponding to only one compound. However, Yuel does not exhibit this memorization characteristic when training on the same dataset. Thus, we speculate that the memorization characteristic is mainly due to the inherent deficiencies of the network architecture. Yuel is equipping with fully-connected (FC) layers that share weights between different compound atoms and protein residues, which we refer to as feature-wise FC layers (see Methods and Figure 3B); other neural networks employ regular FC layers that operate the entire concatenated compound-protein feature matrix, which we refer to as position-wise FC layers (see Methods and Figure 3A). Networks with position-wise FC layers may predict CPI by memorizing the inputs by only optimizing the weights that representing atom-atom interactions in the compound (*W_CC_*) or residue-residue interactions (*W_PP_*) in the protein instead of atom-residue interactions (*W_CP_* and *W_PC_*) (see Methods).

**Figure 2.**
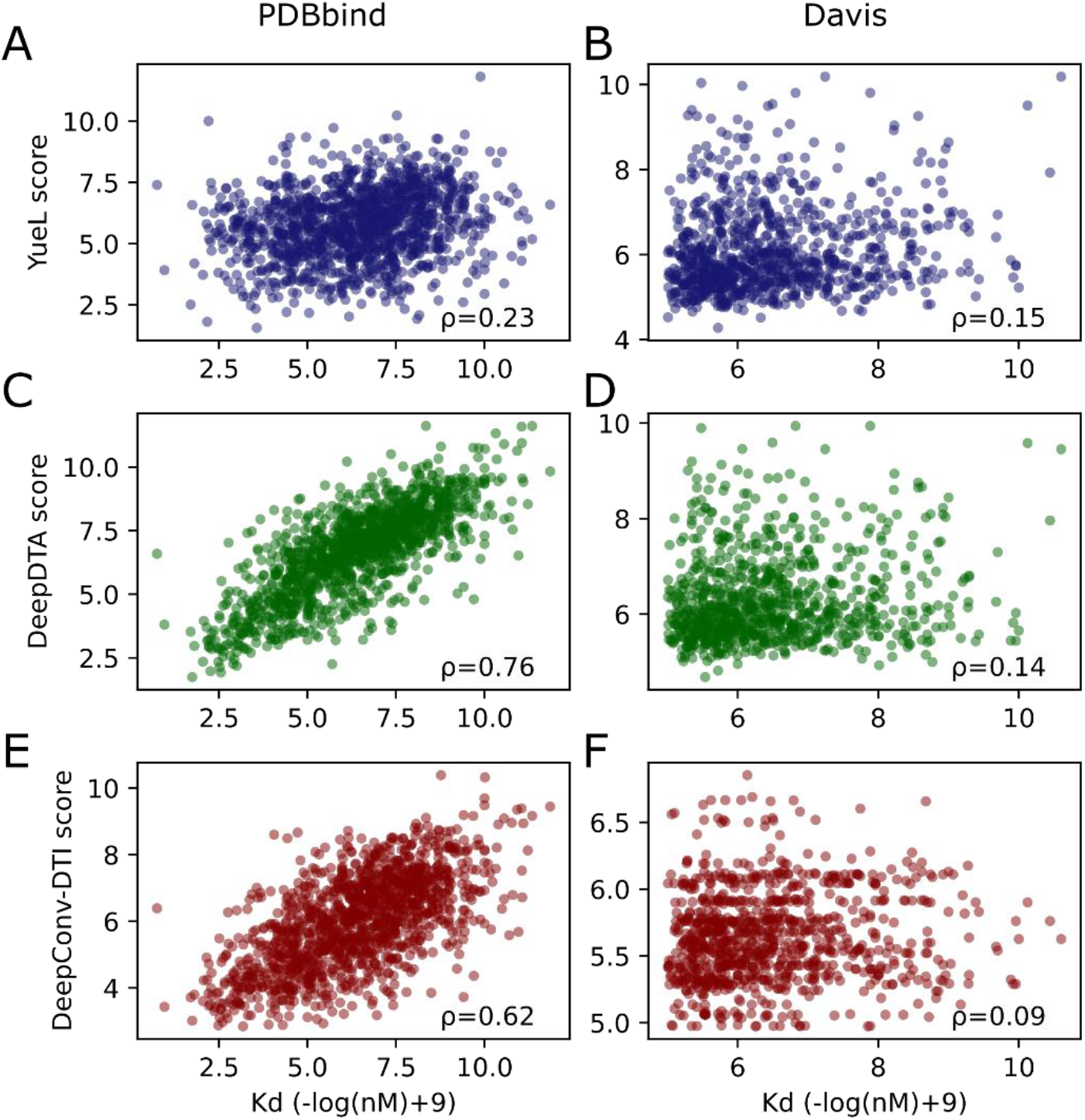
Comparison of Yuel, DeepDTA, and DeepConv-DTI in the shuffled PDBbind and Davis datasets. The models are trained on the normal training set, but the amino acids in each sequence of the test set are shuffled. (A) The relationships between Yuel-predicted affinities and the experimental affinities in PDBbind/test. (B) The relationships between Yuel-predicted affinities and the experimental affinities in Davis/test. (C) The relationships between DeepDTA-predicted affinities and experimental affinities in PDBbind/test. (D) The relationships between DeepDTA-predicted affinities and experimental affinities in Davis/test. (E) The relationships between DeepConv-DTI-predicted affinities and experimental affinities in PDBbind/test. (F) The relationships between DeepConv-DTI-predicted affinities and experimental affinities in Davis/test.

**Figure 3.**
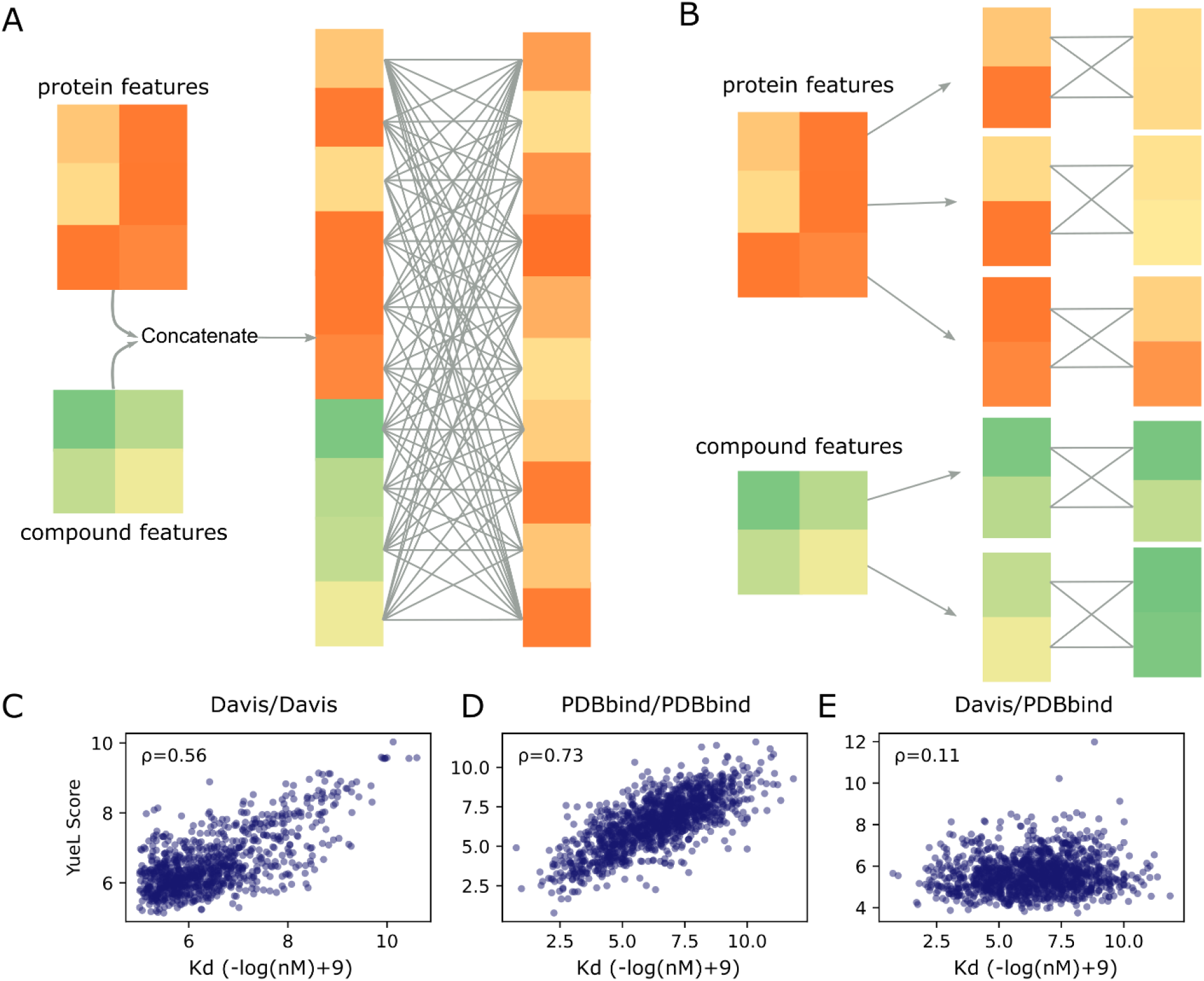
Comparison of position-wise FC layers and feature-wise FC layers. (A) The protein features and the compound features are first concatenated and flattened to a 1-D feature vector. The 1-D feature vector is then subject to full-connected layers. (B) The protein features and the compound features are first split to individual residue features and atom features. Each residue feature and each atom feature is subject to fully-connected layers, individually. Finally, the residue features and the atom features are multiplied to obtain an attention matrix. (C) The affinities predicted by Yuel-cc versus the experimental affinities when training on Davis/train and testing on Davis/test. (D) The affinities predicted by Yuel-cc versus the experimental affinities when training on PDBbind/train and testing on PDBbind/test. (E) The affinities predicted by Yuel-cc versus the experimental affinities when training on Davis/train and testing on PDBbind/test.

To further compare feature-wise FC layers and position-wise FC layers, we create a new version of Yuel by using the position-wise FC layers (Figure S2). We refer to this version of Yuel as Yuel-cc (concatenated version of Yuel). We train and test Yuel-cc on Davis and PDBbind datasets. We find that the Spearman coefficients of Yue-cc are 0.56 and 0.73 in the Davis and PDBbind datasets, respectively (Figure 3C&3D). However, if we train Yue-cc on Davis but test it on PDBbind, the Spearman coefficient is 0.11 (Figure 3E). Thus, by using the position-wise FC layers, Yuel-cc loses the generalizability.

### Identifying hotspot atoms and hotspot residues

When encoding the protein feature matrix, each row corresponds to a residue, and each column corresponds to a specific feature. Similarly, each row of the compound feature matrix corresponds to an atom, and each column corresponds to a feature of the atom. Yuel calculates the outer product of the compound feature matrix by the protein feature matrix to obtain an attention matrix, thus, each element of the attention matrix corresponds to the interaction between a compound atom and a protein residue. Therefore, by comparing the values of different rows in the attention matrix, we can compare the probability of different compound atoms interacting with the protein so as to find the atoms most likely to bind the protein, which we refer to as hotspot atoms; similarly, we refer to residues that are most likely to interact with compounds as hotspot residues. Since different compound atoms have different predicted interactions with different residues (Figure 4A), we adopt a strategy of adding up the interactions of an atom to all residues as the possibility that this atom is a hotspot atom. This strategy results in an average AUC of 0.58 (Figure 4B) for PDBbind. Similarly, the predicted interactions of most hotspot residues are ranked within 30 (Figure 4C). Particularly, for 2IHQ, Yuel can predict 9 out of the 11 hotspot atoms (Figure 4D&4E) and 11 out of the 18 hotspot residues (Figure 4F).

**Figure 4.**
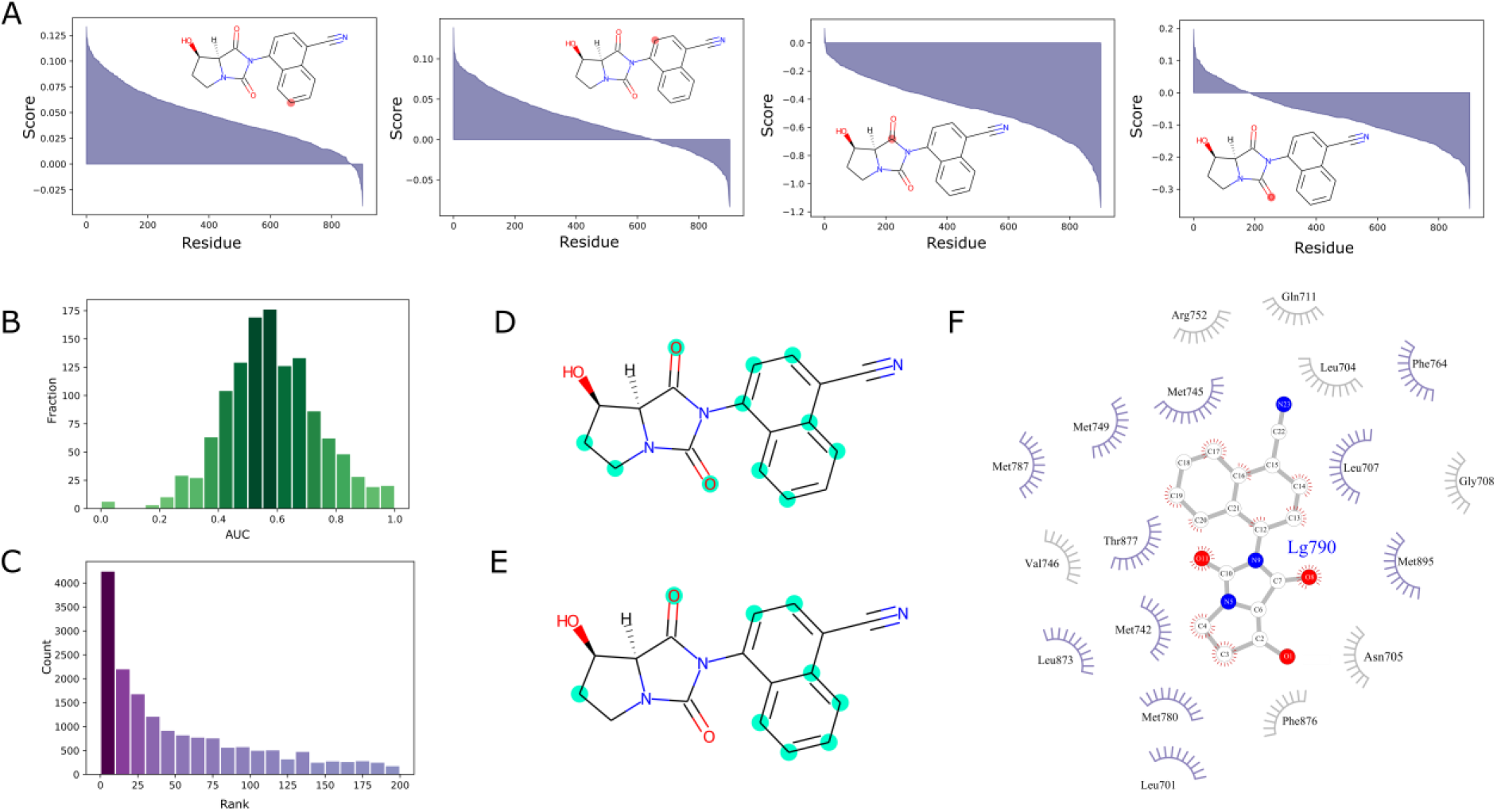
(A) The predicted protein-binding interactions of fours atoms in N-Aryl-Hydroxybicyclohydantoin (LG790), the ligand of the rat androgen receptor (PDB ID: 2IHQ). (B) The histogram of AUC calculated by using Yuel to predict hotspot atoms in LG790. (C) The ranks of hotspot residues in the protein of 2IHQ. (D) The ground-truth hotspot atoms in LG790. (E) The predicted hotspot atoms in LG790. (F) The predicted hotspot residues in the protein of 2IHQ.

## METHODS

### The Architecture of Yuel

*Yuel* requires two inputs to estimate interactions between a specific compound and the protein target: (i) compound’s 2D structure (SMILES^20^ or INCHI^21^ code) and (ii) protein’s amino acid sequence. Given the protein sequence, *Yuel* first encodes it using BLOSUM62 matrix^22^. Each residue corresponds to a column in the BLOSUM62 matrix. The features of non-standard amino acids are zero-initialized. Then, the protein features are updated through three 1-D convolution layers. The sequence is zero-padded to a fixed length of 2048. Given the SMILES of the compound, *Yuel* first employs rdkit^23^ to represent it by a graph (*N*, *V*, *E*), where *N* is the number of nodes, *V* is the feature vector of each atom, and *E* is the feature vector of each bond. The feature of each atom is the concatenation of the one-hot encoding of atom type, number of bonds, bond type, mass, and charge vectors. The feature of each bond is the bond order. The graph is then subject to two graph convolution layers to update the features. The protein features and the compound features are then subject to 5 feature-wise fully-connected layers, separately. Following the fully-connected layers, the protein feature matrix and the compound feature matrix are then multiplied to obtain an attention matrix. Each row of the attention matrix corresponds to a compound atom, and each column corresponds to a protein residue. The maximum value of each column is then selected and added up to get the final predicted CPI. We implement this network using TensorFlow 2.3.1^24^ and Sonnet 2. The GNN is implemented by the *graph_nets^25^* library.

### Position-wise fully-connected layers and feature-wise fully-connected layers

The major difference between Yuel and other neural network models lies in the fully-connected layers. In other neural network models, such as DeepDTA and DeepConv-DTI, the CPI is predicted by a function of the concatenated compound-protein feature matrix (Equation (1)):

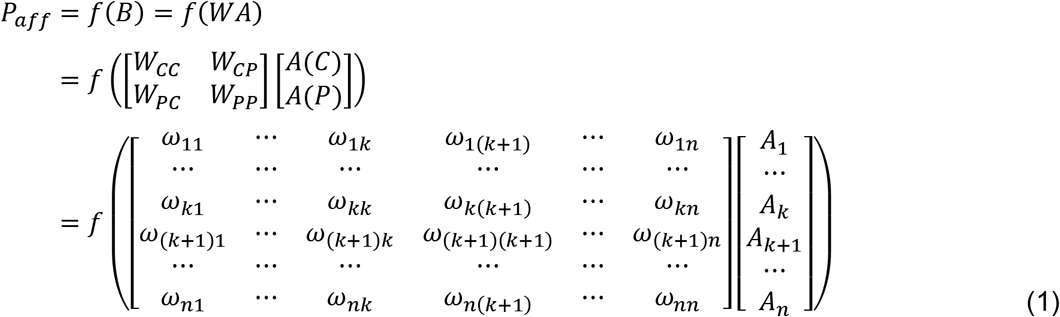

where *B* is the concatenated compound-protein feature matrix;*W* is the weight matrix of the FC layer; *A(C)* is the compound feature matrix before the FC layers; *A(P)* is the protein feature matrix before the FC layer; *A* is the concatenated compound-protein feature matrix before the FC layer; *W_CC_* is the weight matrix for atom-atom interactions in the compound; *W_PP_* is the weight matrix for residue-residue interactions in the protein; *W_CP_* and *W_PC_* are the weight matrices for compound-protein interactions; ω_ij_ is the weight between *A_i_* and *A_j_*, *A_1_*, *A_2_*, ..., and *A_k_* are the features of compound atoms before FC layers; *A_k+1_*, *A_k+2_*, ..., and *A_n_* are the features of protein residues before FC layers. Importantly, the FC layer is applied directly to the concatenated compoundprotein feature matrix, and the weight vector ω_ij_ of the FC layer is dependent on the position *i* and *j* of the compound atom (*i*,*j* ≤ *k*) or protein residue (*i*,*j* > *k*). Thus, we refer to this kind of FC layer as position-wise FC layer (Figure 3A).

In Yuel, the CPI is predicted by a function of the outer product of the compound feature matrix and the protein feature matrix (Equation (2)).

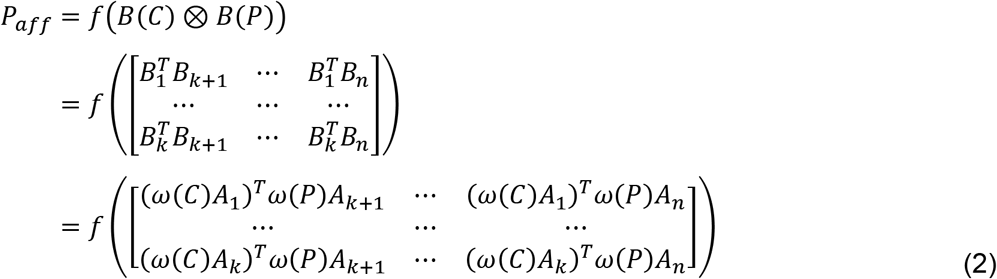

where P_aff_ is the predicted affinity; *B(C)* is the final compound feature matrix; *B(P)* is the final protein feature matrix; *B_1_*, *B_2_*, ..., and *B_k_* are the features of compound atoms; *B_k+1_, B_k+2_, ..., B_n_* are the features of protein residues; *ω*(*C*) is the weight matrix for the compound; *ω*(*P*) is the weight matrix for the protein; the outer product is calculated by treating the compound feature matrix and the protein feature matrix as vectors with each element as a vector of the features of a specific compound atom or protein residue. Crucially, the compound feature matrix *(B(C))* and the protein feature matrix *(B(P))* are processed by FC layers, i.e. *ω*(*C*) and *ω*(*P*), separately. The weight vector *ω*(*C*) of the compound FC layers is not dependent on the position of compound atoms and it is the same for all compound atoms; similarly, the weight vector *ω*(*P*) of the protein FC layers is the same for all protein residues. Thus, we refer to these FC layers as feature-wise FC layers (Figure 3B).

There are three advantages of feature-wise FC layers compared to position-wise FC layers. First, the weights in position-wise FC layers are dependent on the position of atoms and residues. However, when predicting CPI, the protein sequence may lack some residues, which changes the positions of residues, and the position of atoms in the compound may also be different if using a different encoding algorithm. Thus, the position-wise FC layers may result in different affinities for the same compound and the same protein. Second, most parameters of the weight matrix of the position-wise FC layers are focused on atom-atom interactions in the compound (*W_CC_*) and residue-residue interactions in the protein (*W_PP_*), while atom-residue interactions (*W_CP_* and *W_PC_*) are the critical ones for determining CPI, thus, the position-wise FC layers may make the network inclined to memorize the 2D structure of the small molecule and the sequence of the protein by optimizing *W_CC_* and *W_PP_* instead of *W_CP_* and *W_PC_*. Finally, since the weight matrix of the feature-wise FC layers is the same for all atoms or residues, it has far fewer parameters than the weight matrix of the position-wise FC layers, which makes the training of feature-wise FC layers much more efficiency.

### Dateset completeness

We propose a metric, the completeness of a dataset *C_DS_* = *N_interaction_*/(*N_protein_ · N_compound_*), to evaluate whether a dataset is suitable for training a neural network model for CPI prediction. *N_protein_* is the number of proteins; *N_compound_* is the number of compounds; *N_interaction_* is the number of interactions in the dataset. If the completeness of a dataset is 1, it means that the interaction between each compound and each protein is included in the dataset. The completeness of PDBbind, Davis, and Metz is 0.00052, 0.35, and 0.14, respectively. Training DeepDTA or DeepConv-DTI on a dataset with an extremely low completeness (e.g. PDBbind) will make the network inclined to predict CPI by memorizing the protein or the compound structure because the *N_compound_*/*N_interaction_* ratio or the *N_protein_*/*N_interaction_* is close to 1 (Figure 2A&2C&2E). When training on a dataset with a high completeness (e.g. Davis), the overfitting issue in DeepDTA and DeepConv-DTI is alleviated (Figure 2B&2D&2F). Unlike other NN-based methods, the reliability of *Yuel* does not depend on the completeness of the training set.

### Hyper-Parameter Selection

The number of atom features is 82, by using a similar encoding method in Shuya Li’s work^13^. The first 63 atom features is the one-hot encoding of the atom element in the list of C, N, O, S, F, Si, P, Cl, Br, Mg, Na, Ca, Fe, As, Al, I, B, V, K, Tl, Yb, Sb, Sn, Ag, Pd, Co, Se, Ti, Zn, H, Li, Ge, Cu, Au, Ni, Cd, In, Mn, Zr, Cr, Pt, Hg, Pb, W, Ru, Nb, Re, Te, Rh, Tc, Ba, Bi, Hf, Mo, U, Sm, Os, Ir, Ce, Gd, Ga, Cs, and “unknown”; 6 features are the one-hot encoding of the degree of the atom; 6 features are the one-hot encoding of the explicit valence of the atom; 6 features are the one-hot encoding of the implicit valence of the atom; the last feature is whether the atom is aromatic. The number of bond features is 6, indicating whether the bond is single, double, triple, aromatic, conjugated, or in a ring. The number of residue features is 20, the number of amino acid types. The number of GNN layers is 2 and the number of CNN layers is 3. We choose 2 as the number of GNN layers because we consider each atom group as the center atom, the neighbor atoms of the center atom, and the neighbor atoms of the neighbor atoms. We choose 3 as the number of CNN layers such that each residue fragment contains 7 residues. After the last CNN layer, we concatenate 4 protein feature matrices together: the protein feature matrix before the first CNN layer, the protein feature matrix after the first CNN layer, the protein feature matrix after the second CNN layer, and the protein feature matrix after the third CNN layer. The max length of compound vector is 64 and the max length of protein vector is 2048. Since our model only uses the heavy atoms in compounds, 64 as the number of atoms is enough to cover most compounds; similarly, 2048 as the number of residues is enough to cover most proteins. The size of the feature dimension is 32 for most layers in the model. Before multiplying the compound feature matrix and the protein feature matrix, we subject them to 5 fully-connected layers, separately. The size of the feature dimension in the first 4 linear layers is 32*8, and the size of the last linear layer is 32*50.

### Training

The objective is to minimize the mean squared error of the predicted binding affinity and the experimental binding affinity. During the training process, we use a mini-batch of 8 and the Adam stochastic optimizer^26^ to optimize the model parameters. For each training batch, compounds with different numbers of atoms and proteins with different numbers of residues are zero-padded to obtain the same input feature lengths. During the training process, the padded regions of features are masked so that they do not contribute to the calculation of the losses and gradients. Yuel has about two million learnable parameters. A single Yuel model has 1411523 parameters if the *“nfeat”* variable is 32. We also train Yuel by setting the “nfeat” variable as 8, and the performance does not change significantly (Figure S3). Training Yuel by 200 epochs on a dataset containing 440000 CPIs needs 34 hours on a Linux server with one NVIDIA Tesla T4 GPU.

### Datasets

Our benchmark datasets include the PDBbind dataset^18^, the Davis dataset^17^, and the Metz dataset^19^. The PDBbind dataset is a high-quality set of protein-ligand complexes with available structural data and corresponding binding affinities. The PDBbind dataset is compiled from the PDBbind database (version 2018, the general set), which contains a high-quality set of protein-ligand complexes with available structural data and corresponding binding affinities. Each complex was provided with an affinity value of certain measurement type. The Davis dataset contains interactions of 72 kinase inhibitors with 442 kinases covering >80% of the human catalytic protein kinome. The Davis dataset is obtained from the Supplementary Information of Davis et al. work^17^. The Metz dataset is obtained from the Supplementary Information of Metz et al. work^19^. The PDBbind dataset contains 2509 small molecules and 10251 proteins, with a total of 13,311 interactions. On average, each small molecule corresponds to only 5.3 interactions and each protein corresponds to only 1.3 interactions. The Davis dataset contains 68 small molecules and 379 proteins, with a total of 9125 interactions. On average, each small molecule corresponds to 134.2 interactions, and each protein corresponds to 24.1 interactions. The Metz dataset contains 1471 small molecules and 172 proteins, with a total of 35307 interactions. On average, each small molecule corresponds to 24.0 interactions, and each protein corresponds to 207.7 interactions. We divide each dataset into a training set and a test set at a ratio of 8 to 2.

## DATA AVAILABILITY

Source codes and test data are deposited in: *https://bitbucket.org/dokhlab/yuel*. We provide a website to use Yuel: *https://yuel.dokhlab.org* or *https://dokhlab.med.psu.edu/cpi*.

## ACKNOWLEDGEMENTS

We acknowledge support from the National Institutes for Health (R35 GM134864), National Science Foundation (2040667) and the Passan Foundation. We appreciate Congzhou Sha for his enthusiastic proofread of the manuscript.

