## Supplemental Information for "*Yuel*: Compound-Protein Interaction Prediction with High Generalizability"

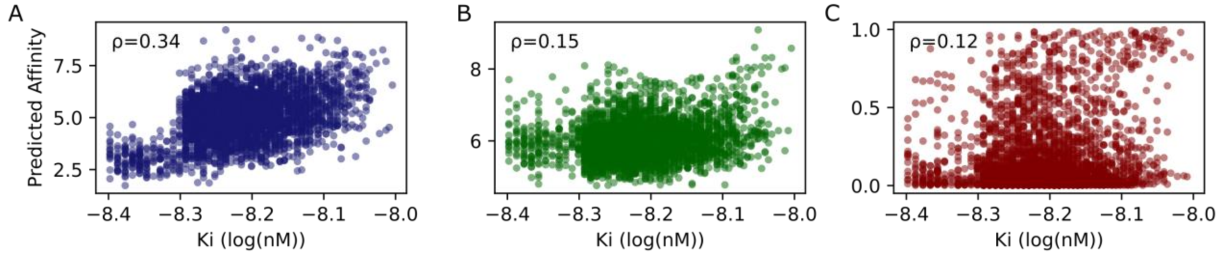


**Figure S1. Comparison of the ability of Yuel, DeepDTA, and DeepConv-DTI to predict compound-protein interactions in the Metz dataset.** All models are trained on the Davis dataset. (A) The relationships between the compound-protein interaction predicted by Yuel and the experimental affinity data of the Metz test set. (B) The relationships between the compound-protein interaction predicted by DeepDTA and the experimental affinity data of the Metz test set. (C) The relationships between the compound-protein interaction predicted by DeepConv-DTI and the experimental affinity data of the Metz test set.

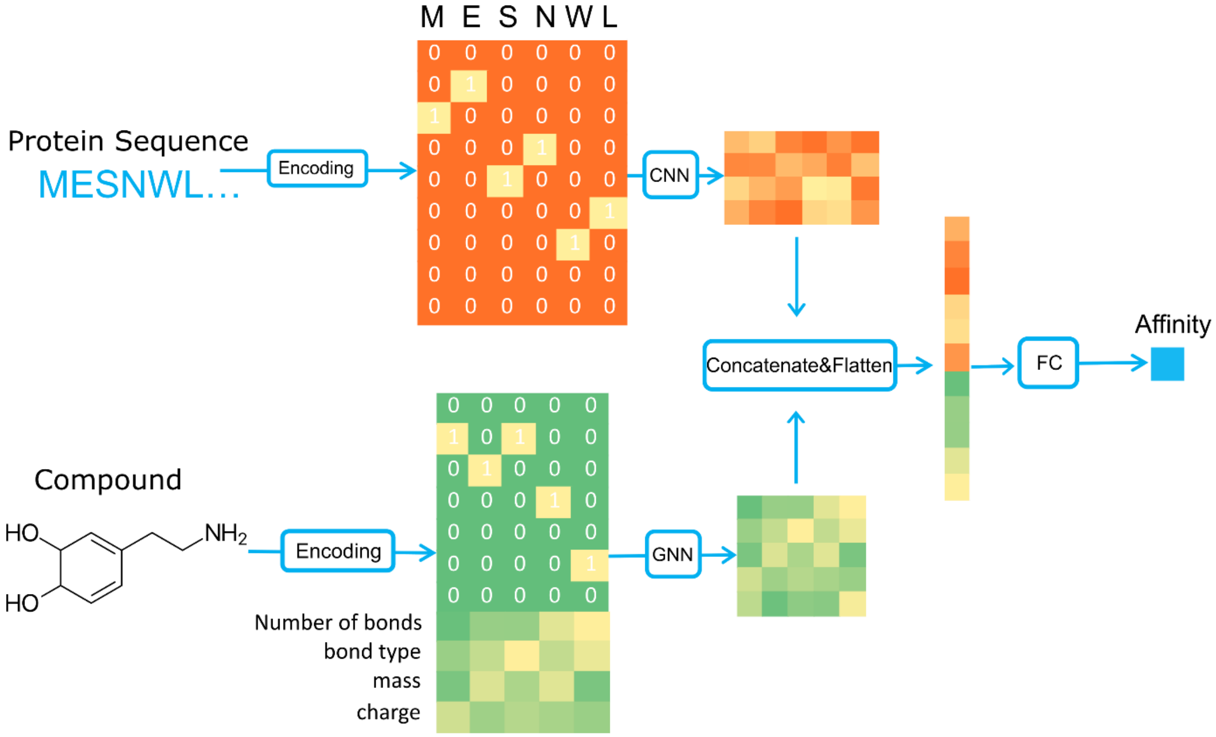


**Figure S2. The architecture of Yuel-cc.** Yuel-cc is the modified version of Yuel by conforming to the encoding-FC paradigm and using the position-wise FC layers.

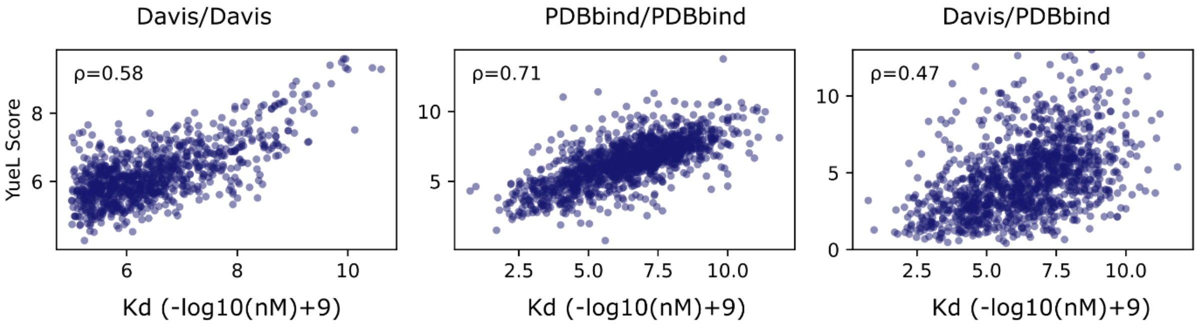


**Figure S3. Test of the performance of Yuel when the number of features is 8.**
